# Exploitation by cheaters facilitates the preservation of essential public goods in microbial communities

**DOI:** 10.1101/040964

**Authors:** Clara Moreno-Fenoll, Matteo Cavaliere, Esteban Martínez-García, Juan F. Poyatos

**Affiliations:** Logic of Genomics Systems Lab (CNB-CSIC), Madrid, Spain; University of Edinburgh, Edinburgh, UK; Molecular Environmental Microbiology Lab (CNB-CSIC), Madrid, Spain

## Abstract

How are public goods^1-4^ maintained in bacterial cooperative populations? The presence of these compounds is usually threatened by the rise of cheaters that do not contribute but just exploit the common resource^5,6^. Minimizing cheater invasions appears then as a necessary maintenance mechanism^7,8^. However, that invasions can instead add to the persistence of cooperation is a prospect that has yet remained largely unexplored^6^. Here, we show that the detrimental consequences of cheaters can actually preserve public goods, at the cost of recurrent collapses and revivals of the population. The result is made possible by the interplay between spatial constraints and the essentiality of the shared resource. We validate this counter-intuitive effect by carefully combining theory and experiment, with the engineering of an explicit synthetic community in which the public compound allows survival to a bactericidal stress. Notably, the characterization of the experimental system identifies additional factors that can matter, like the impact of the lag phase on the tolerance to stress, or the appearance of spontaneous mutants. Our work emphasizes the unanticipated consequences of the eco-evolutionary feedbacks that emerge in microbial communities relying on essential public goods to function, feedbacks that reveal fundamental for the adaptive change of ecosystems at all scales.

## Main text

The threat of cheaters represents at a microbial scale a well-known public good (PG) dilemma, known as the “tragedy of the commons”^9^, and can fundamentally interfere with the sustainability of microbial communities. The necessity of recognizing the consequences of social dilemmas in microorganisms thus becomes essential, given their impact in many aspects of life on Earth, and also its particular relevance to humans in matters of health (microbiome)^10^, and industry (bioremediation, biofuels, etc)^11^. We considered specifically a scenario in which a community is organized as a dynamical metapopulation (i.e., the community is transiently separated into groups)^12^, and the action of a PG is essential for its survival. Spatial structure is a well-known universal mechanism to promote cooperation^13^, which frequently emerges in bacterial populations, for instance, due to the restricted range of microbial interactions^14,15^. However, it is much less understood how the presence of structure affects the maintenance of cooperation when combined with explicit population dynamics (earlier work usually assumed constant population and only examined evolutionary dynamics)^16^. The change in population size associated to the essentiality of the PG can indeed bring about complex eco-evolutionary feedbacks^17-20^, in which both population density and frequency of “cooperators” influence each other. The connection between these feedbacks and spatial structure remains thus an open problem that has started to be addressed only recently^21-22^. We show in this work how such connection can direct to the unforeseen consequence that cheater invasions eventually support cooperation.

To analyze this scenario, we first introduced a stylized *in silico* model considering an initial finite population of agents –representing bacteria– with a given frequency of cooperators (producers of a PG, with a fitness cost) and cheaters (nonproducers, that could have emerged originally from the cooperators by mutation)(Methods). The population is temporarily organized in groups, where interactions take place (fig. S1). These interactions are modelled by means of a PG game with individual reproduction being set by the game payoff ^17,18^. Figure 1A displays a representative trajectory of the model: an increase of the cheating strain, due to its fitness advantage, causes a decrease in population density (less PG available). The demographic fall originates in the end variation in the composition of the groups, facilitating population assortment and the appearance of pure cooperator/cheater groups. Since the groups uniquely constituted by cooperators grow larger, they can ultimately reactivate the global population promoting again new cheater invasions. The whole process manifests in this way as a continuous cycle of decay and recovery of the community (Figs. 1A-B) (Methods). Demographic collapses consequently turn into an endogenous ecological mechanism that causes the required intergroup diversity, supporting the overall increase of cooperators by means of a statistical phenomenon known as Simpson’s paradox^12,22^. To underline that an endogenous ecological process naturally induces the variance that finally rescues cooperation, we introduced the notion of “ecological Simpson’s paradox”.

We then tested these ideas experimentally, by engineering a synthetic PG interaction that is essential for the survival of a microbial population to a bactericidal antibiotic (Fig. 1C). Specifically, we constructed an experimental system in which a synthetic *Escherichia coli* strain (the cooperator/producer) constitutively secretes an autoinducer molecule acting as PG. This molecule is part of a *quorum-sensing* (QS) system foreign to *E. coli*, which includes a cognate transcriptional regulator. We connected this machinery to the expression of a gene that enables the synthetic strain to tolerate the bactericidal antibiotic gentamicin (gm) (fig. S2, the system is a variation of an earlier one^22^)(Methods). A second strain (the cheater/nonproducer) that only utilizes the PG can also be part of the community (we labelled the cooperative and cheater strains with a green and red fluorescent protein, respectively, to make possible population measures) (Methods). Two crucial aspects distinguish in this way the designed setup. First, the presence of PG becomes an essential requirement to tolerate stress (Fig. 1D) (fig. S3). Second, the system exhibits an intrinsic vulnerability, as cheaters could overtake the entire community since they evade the cost of producing the PG (Fig. 1E) (fig. S4). While in this case the presence of cheaters is part of the synthetic design, their emergence as result of mutations is well documented in natural settings^5,6^.

**Fig. 1.**
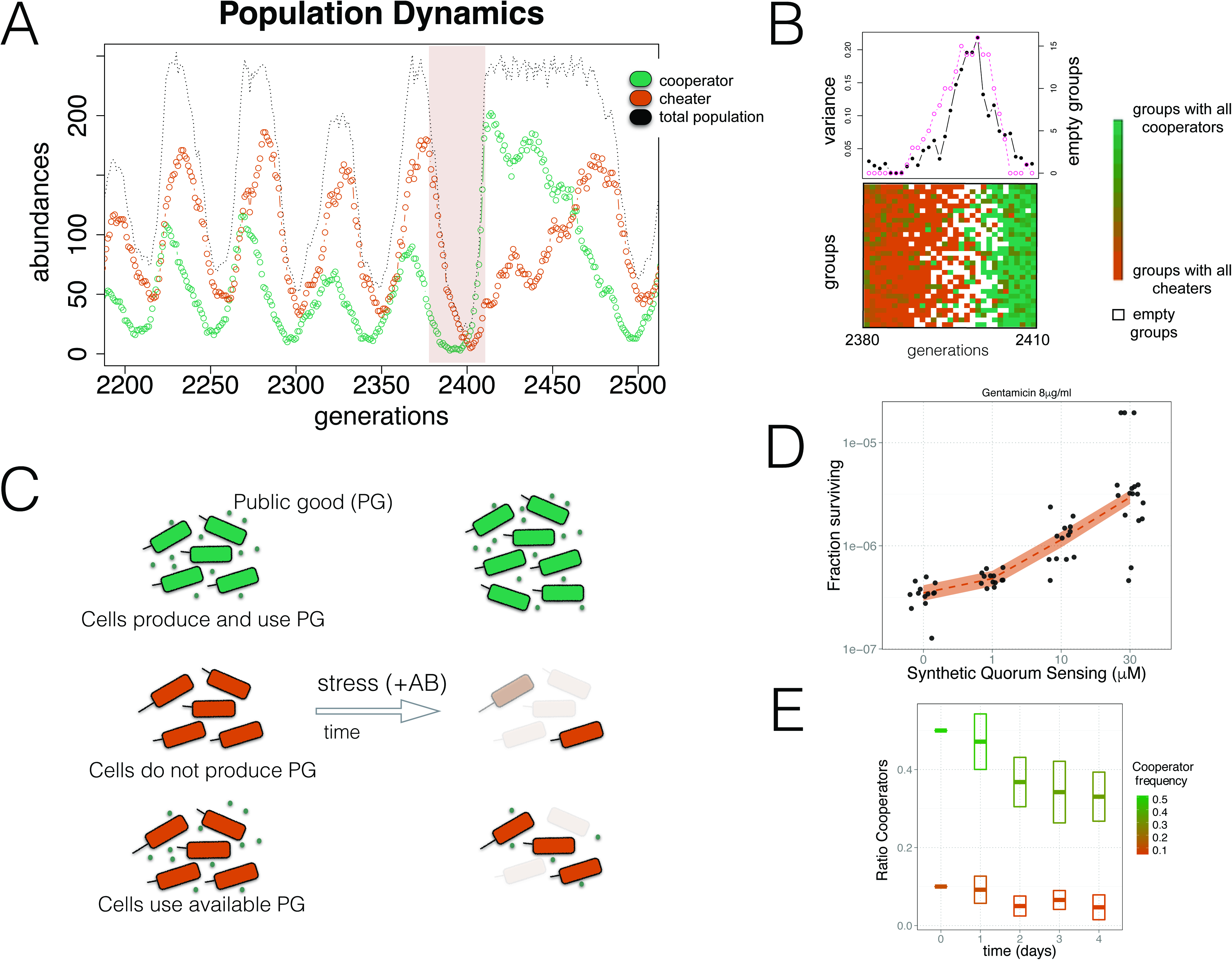
Exploitation by cheaters contributes to the maintenance of cooperation. (**A**) Typical population dynamics obtained with an *in silico* model of a microbial community, which is organized as a transient metapopulation (fig. S1) (Methods). Growth depends on an essential public good (PG) produced by the “cooperator” individuals. When “cheaters” invade, the decline in the amount of PG drives the collapse of the total population. This collapse paradoxically determines its subsequent revival. (**B**) Revival is coupled to the endogenous emergence of variability in the composition of the groups (constituting the metapopulation) when the population is falling^17,18^, and the following occurence of groups only constituted by cooperators (ecological Simpson’s paradox). For the time window highlighted in (A), we display group composition (bottom; ratio of cooperators on each group is colored according to the gradient shown; white squares denote empty groups, of a total of *N*=30), intergroup diversity (top; black curve, quantified as variance in group composition), and number of empty groups (top; pink curve). (**C**) To test experimentally these ideas, we engineered synthetic strains that contribute (green), or not (red), to the production *of quorum-sensing* (QS) molecules acting as an essential PG (fig. S2). A cell community that accumulates (top) or obtains (bottom) the PG in the environment is able to tolerate a bactericidal antibiotic stress (gentamicin, gm), although the latter can survive least because the initial PG is spent. Moreover, a community growing without PG would eventually collapse (middle). (**D**) Survival to gm of a population constituted only by cheaters increases when the essential QS molecules are added to the preincubation medium, before growing under the stress (8μg/ml gm, *N=17* replicas per QS dosage, solid color represent 95% confidence interval of a local polynomial regression). (**E**) Fraction of cooperators in a mixed population decays with time due to the invasion of cheaters, which do not pay the cost of making the PG (fig. S4). This is independent of the initial fraction (10% or 50% of cooperators). Boxes indicate standard deviation linked to the experimental estimation of ratios. See (Methods) for further details and protocols.

The validation of the presented ecological Simpson’s paradox is done using a minimal experimental protocol integrating the most important features of the model: the demographic collapse induced by cheaters, and the subsequent recovery of cooperation supported by the spatial constraints, i.e., the metapopulation. Specifically, the protocol includes two growing periods of PG accumulation and later exposure to stress (gm). Densities of cooperators and cheaters are quantified by plating (Methods). Both the *in silico* model and the main attributes of the experimental system predict that a community with different frequency of cooperative and noncooperative strains would accumulate a distinct amount of PG, and thus present different tolerance to stress. To confirm this, we engineered an initial population with density ~10^4^cells/ml and different composition (fig. S5)(Methods). Figure 2A confirms the differential tolerance to stress (inset illustrates the resultant range of accumulation of QS molecules), whereas Fig. 2B additionally shows the effect on tolerance of rising the strength of stress for a fixed composition (20% cooperators). Note that the fraction of producers in these final populations does not vary substantially (fig S6).

**Fig. 2.**
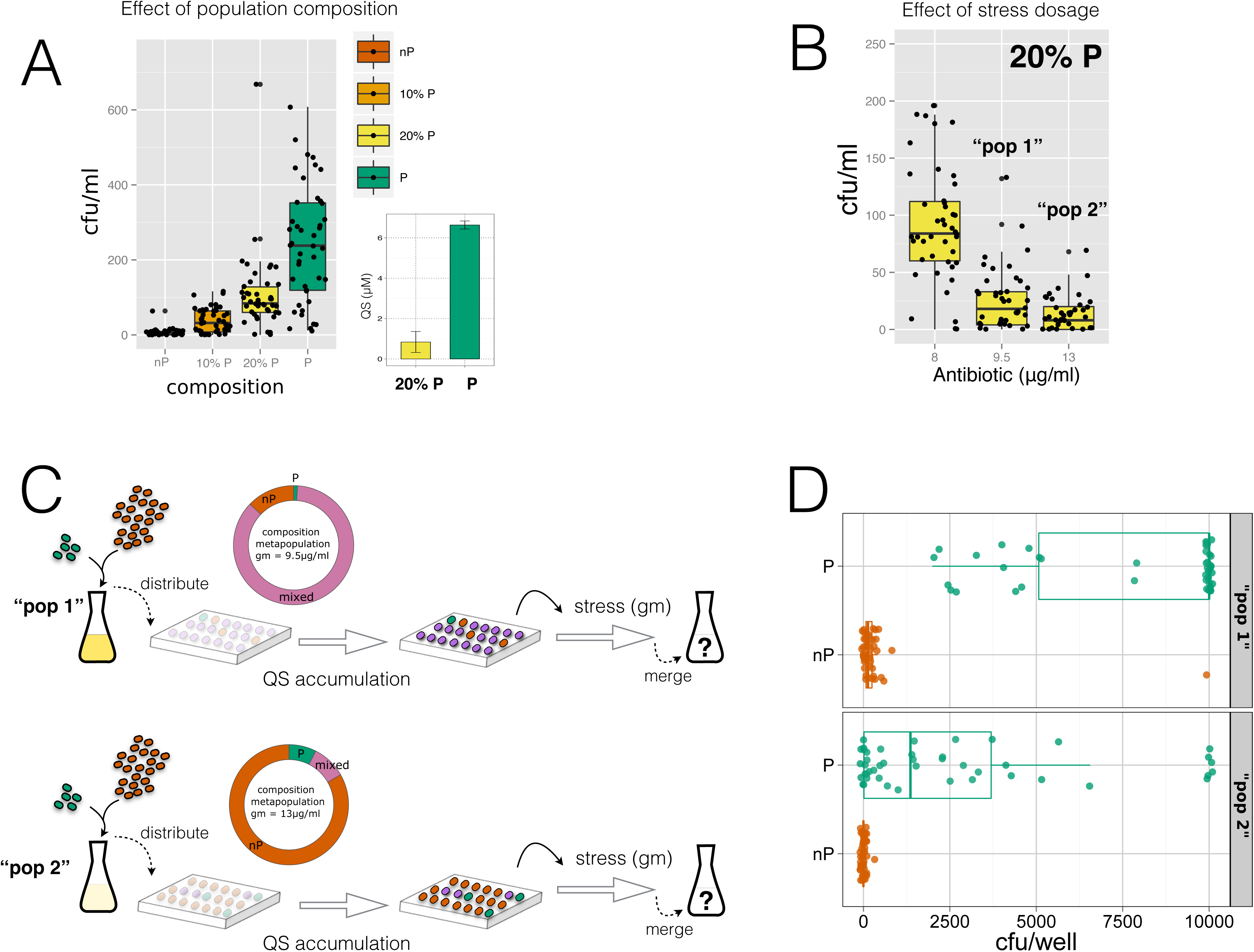
Experimental test of the ecological Simpson’s paradox. (**A**) Survival population to gentamicin (gm) exposure from an initial density of ~10^4^ cells/ml and different population composition. Cells grow for 15.5 hrs (T_1_) without stress to be then 1/10 reseeded in a medium with 8μg/ml of gm, in which they grow 8.5 hrs (T_2_) (Methods). Each dot represents a single replica (*N*=45), and box plots represent associated statistical parameters. Cooperators/cheaters labelled as producers (P)/nonproducers(nP) of the public good, respectively. (Right inset) Accumulation of *quorum-sensing* molecules (QS) at T_1_; population constituted by 20% or all Ps (Methods). (**B**) Survival population to three gm dosages. Initial density of ~10^4^ cells/ml, with a composition of 20% Ps. Dots and box plots as in (A), *N*=45. Population densities, and associated dosages of 9.5μg/ml and 13μg/ml, are correspondingly termed as “pop 1” and “pop 2” conditions. The frequency of P remains approximately the same at the end of the protocol (fig. S6). (**C**) Cartoon of a second round of the experimental protocol. Initial conditions correspond to those of either “pop 1” or “pop2” in (B). Each population is first distributed in a multiwell plate representing a metapopulation, followed by a period of accumulation of QS (T_1_) and stress (T_2_), as before (low initial densities on each well are represented here by weak opacity). The population is then quantified by plating and considered as a whole (merged metapopulation). Note the enrichment of groups of nPs for the “pop 2” experimental condition, and also the appearance of some groups constituted by only Ps. (**D**) Groups of Ps tolerate better the stress than groups of nPs, and lead to the recovery of the cooperation. We quantified the characteristic tolerance by engineering replica populations of only Ps and only nPs with the “pop 1” and “pop 2” associated cell densities and gm dosages, and then measured recovery of each well (each dot represents the fate of a replica well, *N*=45). A maximal value of ~10000 cfu/well denotes very strong recovery. Mixed wells with low initial density normally exhibit the fate of nPs at these high dosages (fig. S8). Note that nPs show almost no recovery in the “pop 2” conditions (bottom, gm = 13μg/ml). See also main text.

Next, by distributing the initial full population into a metapopulation structure –before the rounds of QS accumulation and stress– we showed experimentally the basic features of the ecological Simpson’s paradox. In particular, we selected as initial conditions the ones obtained after a large mixed population experienced a first round of comparatively strong stress (Fig. 2B, outcomes of an earlier round with a initial density of ~ 10^4^cells/ml constituted by 20% producers). As in the model, the reduced accumulation of PG in the mixed population causes a demographic collapse, which depends on how essential the PG becomes (the strength of stress). We considered explicitly the resulting populations after experiencing a medium (9.5μg/ml) and strong (13μg/ml) gm stress. These corresponded on average to ~24cfu/ml, and ~8cfu/ml, respectively (in what follows, we labelled them as “pop 1” and “pop 2”; the frequency of cooperators remains ~20%). When we subsequently implemented a second round of the protocol with these initial population conditions, and the corresponding (medium and strong) gm dosages, we obtained two drastically different initial metapopulation distributions (Fig. 2C) (fig. S7). Notably, a low-density population (“pop 2”) generates groups with very few cells and assortment of cooperators, which is the group class that best tolerate stress (the characteristic behavior of this class is shown in Fig. 2D). Moreover, mixed groups and those composed by only cheaters do not grow, on average, with strong gm dosage (Fig. 2D) (fig. S8). In this way, when all the groups of the metapopulation are pooled together, cooperators typically increase in frequency and population density recovers, in what is a manifestation of the ecological Simpson’s paradox previously described (fig. S8) (Methods). Hence, the demographic decay of the population caused by the invasion of cheaters reveals crucial to attain the necessary heterogeneity among the various groups, and allows the recovery of cooperation. For a large population, groups exhibit a similar composition, on average, and as a result the behavior in the occurrence of the metapopulation is the same as in the absence of any structure, i.e., every well is in effect a replica of the same condition.

Two additional constraints could modify the dynamics above, which associate to general aspects of resistance and tolerance to antibiotic stress^23^. The first one identifies a possible trade-off between the build up of PG and the lag time of bacteria before exposure to the antibiotic. As Fig. 2A illustrated, the more PG a population accumulates (i.e., the longer it grows), the more it tolerates stress. However, when extensive growth implies a deterioration of the environmental conditions, PG could lose its functional activity^24^ (Fig. 3A)(fig. S9). Growing too much could be problematic. Moreover, when a bacterial population stays some time in saturating conditions, it presents a longer lag phase that indirectly protects bacteria from antibiotics^23^, independently on whether PG is available or not (Fig. 3A). Within the previous context of the recovery due to the ecological Simpson’s paradox (Fig. 2C-D), low initial densities assure a regime further from saturation at the end of the accumulation period. This implies less decay of the PG and shorter lag, i.e., revival is under these conditions mostly linked to the action of the PG. The second constraint relates to the spontaneous emergence of mutants, which can resist the antibiotic and thus enable recovery of cheater populations in the absence of PG. This type of rescue is more difficult as the antibiotic dosage is increased, e.g.,^25^ (fig. S10), and also underlines a constraint on the accumulation time of PG (the longer this period, the bigger the population and the chance for a mutant to arise) (Fig. 3B-C). Of note, both of these restrictions are less significant when the collapse of the population is very strong (“pop 2” situation), that is when the best conditions exist for the ecological Simpson’s paradox.

**Fig. 3.**
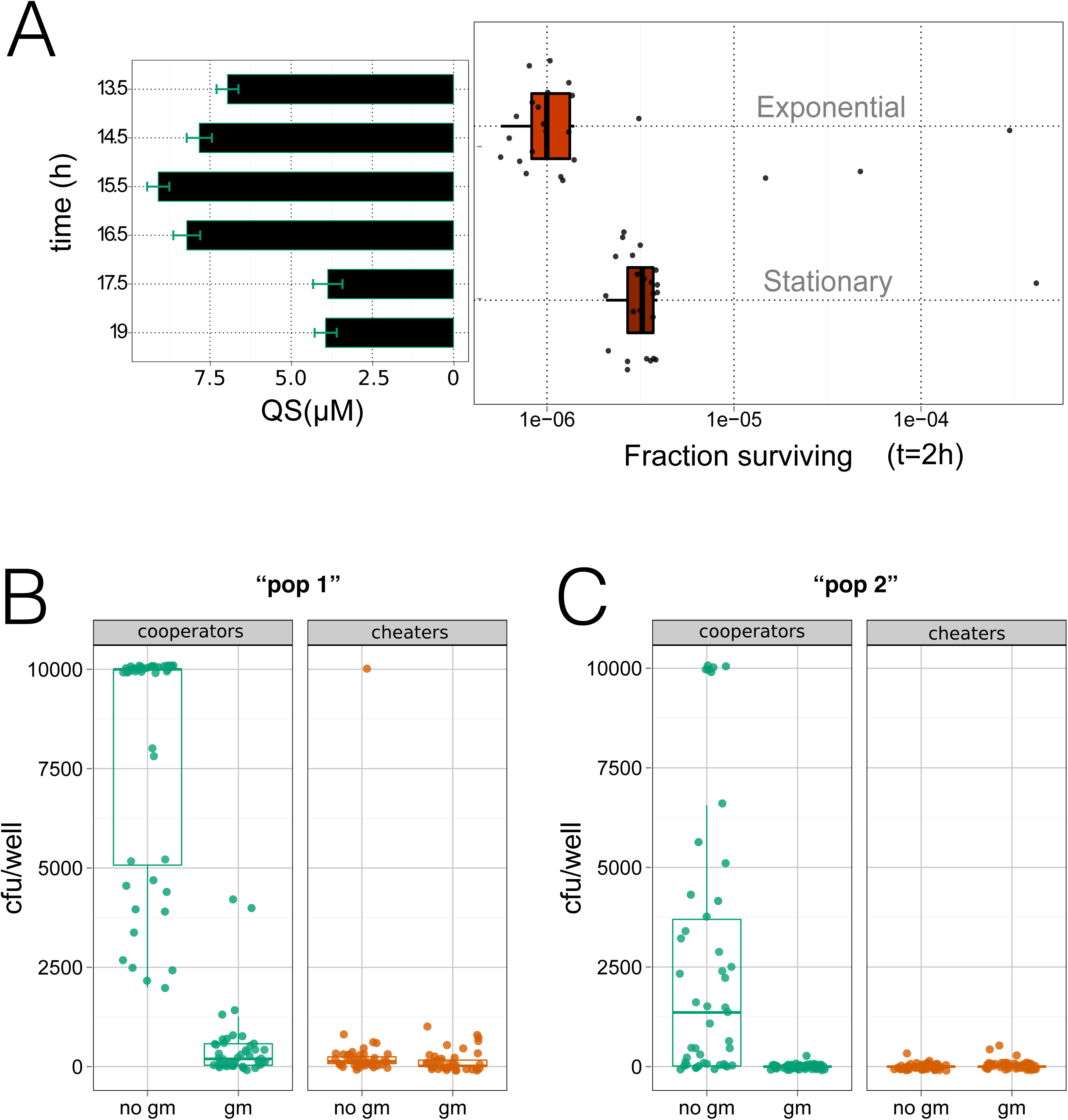
Additional constraints influence the maintenance of public good-based communities under antibiotic stress. (**A**) Cheater strains in exponential or saturating phase were resuspended in medium with gentamicin (gm) (Methods). (Right) Cells in exponential phase experienced less antibiotic tolerance^23^ (gm =12μg/ml, dots correspond to *N*=45 replicas, box plots indicate associated statistical parameters). (Left) Concentration of *quorum-sensing* (QS) molecules decays as a function of time of growth^24^ (figure S9), starting from an initial low-density of cooperators, i.e., “pop 2” density condition in Fig. 2. Bars represent measurement errors associated to QS estimation (Methods). Starting from these “pop 2” initial densities, and after an accumulation time of 15.5 hrs (T_1_), cells are in exponential phase, and the amount of QS is maximal. Recovery is thus strongly linked to the presence of the public good (PG). (**B-C**) Emergence of spontaneous mutants to gm. Initial populations of cooperators and cheaters are subjected to an accumulation and stress protocol under the “pop 1” and “pop 2” conditions (same as Fig. 2; black dots represent replicas, *N=45;* box plots represent statistical parameters, color codes as Fig. 2). We repeated the experiment for each strain and dosage, so that one can quantify the typical resulting population, and also the mutant subpopulation (by plating with –gm– and without –no gm– antibiotic; the specific plating dosage corresponds to that of the matching growing conditions). Emergence of spontaneous mutants is reduced at higher dosage (C). Tolerance is most significantly associated in this regime to the presence of the PG.

This study emphasizes that the combination of spatial constraints and specific attributes of the PG (e.g., essentiality) can be crucial for the outcome of the eco-evolutionary dynamics in cooperative bacterial communities. This is important in circumstances where the PG aids tolerance to stress, when nonintuitive effects may appear: an increase in cheaters frequency, or AB dosage, can actually precede the recovery of cooperation in the population. The particular experimental setup also provides synthetic strains with essential public goods; an original tool to test the demographic risks of cheaters invasions and implement experimentally a microbial tragedy of commons^9^, where cooperation is tied to population survival. Our results show overall that in this framework, spatial constraints, growth phase and the emergence of noninteracting mutant clones are aspects that should be considered to fully appreciate the resilience of social interactions in bacterial populations. In addition, synthetic communities emerge as tractable experimental models in which to begin to understand the tight ecological and evolutionary feedbacks increasingly observed in ecosystems worldwide^26^.

## Supplementary Information

Figures S1-S10

Table S1

## Acknowledgements

We would like to thank John Chuang for strains, and A. Couce, R. Díaz-Uriarte, D. Bajic, A. Sánchez, A. Pascual, O. X. Cordero, and V. de Lorenzo for helpful discussions. This work was partially supported by grants from La Caixa Foundation PhD fellowships, the Spanish Ministry of Economy and Competitiveness, and the British Engineering and Physical Sciences Research Council.

## Methods

### Ecological public good model

We used a model first described in 21to simulate the dynamics of a population whose growth is based on an essential public good (PG). It is based on a one-shot PG game27 in which agents can contribute (“cooperators”) or not (“cheaters”) to the PG in groups of size *N.* Contributing implies a cost *c* to the agents. Group contributions are then summed, multiplied by a reward factor *r* (that determines the efficiency of the investments and the attractiveness of the PG) and redistributed to all group members, *irrespectively* of their contribution. The PG game is characterized by the parameters *N, r* and *c* (group size, efficiency and cost of the PG, respectively, where we fixed *c* = 1 without loss of generality).

Every simulation starts with an initial population constituted by a common pool of *k* identical agents in the *cooperator* state, where *k* is the maximal population size (carrying capacity), to be updated in a *sequential* way as follows (see also Figure S1):

i. The common pool is divided in randomly formed groups of size *N* (i.e. *N* is the total number of individuals and empty spaces in each group). The number of formed groups is then ⌊*k/N*⌋.
ii. In each one of the (non-empty) groups, a one-shot PG game is played. This means that cheaters receive the payoff P_cheater_ = *icr/(i + j)*, while cooperators receive the same payoff minus a cost, i.e. P_cooperator_ = P_cheater_ − *c*; with *i,j* being the number of cooperators and cheaters in the group, respectively, and *i + j ≤ N.* After the interaction the grouping of individuals is dissolved.
iii. Each individual can replicate (duplicate) with a probability that is calculated by dividing its payoff by the maximal possible one (i.e. the payoff obtained by a cheater in a group of *N*−1 producers). Each cooperator that replicates generates an offspring that is either a cheater (with probability *v*) or an identical cooperator (with probability 1−*v*).
iv. Individuals are removed with probability *δ* (individual death rate).

In simpler words, the life cycle of the computational model is characterized by two distinct *stages.* In stage I (steps i–ii), the population is structured in evenly sized randomly formed groups in which the PG game is played. In stage II (steps iii-iv, after groups disappear), each individual replicates according to the group composition (and payoff) experienced in stage I. Replication can happen only when the current total population is less than the maximal population size, *k*, i.e. there exits empty space (empty spaces are calculated by considering *k* minus the current amount of individuals in the population). If more individuals could replicate than the available empty space, only a random subset of them ultimately replicates (of size the number of empty spaces available).

### Media, growth conditions and chemicals

Strains were grown in LB broth (10gl1 casein peptone, 5gl-1 yeast extract, and 5gl-1 NaCl) at 30°Cwith constant shaking. Overnight cultures were grown aerobically in flasks (170rpm) at 30°C, and experiments were performed in 96-well plates (Thermo Scientific, Denmark), with 200μl of medium, 50μl of mineral oil (Sigma-Aldrich, MO, USA), and shaking at 10000rpm and 30°C. Where indicated, antibiotics were added to the liquid medium or plate at final concentrations: kanamycin (Km) 50μg/ml, spectinomycin (Sp) 50μg/ml in the construction of strains and 25μg/ml in experiments, and/or gentamicin (gm) with concentration as noted in the experiments. The synthetic *quorum-sensing* molecule N-butyryl-L-homoserine lactone (C4-HSL) was purchased from Cayman Chemical (Mi, USA). Cell dilutions were done in PBS (pH 7.4, 80.6mM Na2HPO4, 19.4mM KH2PO4, 27mM KCl, 1.37M NaCl at 10X, USB Corporation, OH, USA).

### Measurement of population size

Cultures were spread onto 1.5% (w/v) agar plates with five 3mm glass beads for 30s, and are incubated at 30°Cfor 48h (or otherwise indicated). Then, to quantify the cell number of a population we counted colony forming units (CFU) under blue light illumination (LED transilluminator, Safe Imager™ 2.0, Invitrogen, Waltham MA USA). The OD_600_ of cultures was measured in a VICTOR2x 2030 Multilabel Reader machine (Perkin Elmer, Waltham, MA, USA) with intermittent orbital shaking. For experiments with P and nP strains, Km and Sp were added to the media.

### General DNA techniques, plasmids and strain constructions

The different *E. coli* strains were derivatives of JC1080 (BW25113 *Δsdi∷FRT*)^22^. Oligonucleotides used in this work are indicated in Table S1. Plasmid DNA was prepared using the QIAprep Spin Miniprep kit (Qiagen, Inc., Valencia, CA). When required DNA was purified using the NucleoSpin Extract II (Macherey-Nagel, Düren, Germany). Colony PCR was performed by transferring cells directly from fresh agar plates into PCR reaction tubes.

The pZS4int-rhIL-GmLAA plasmid was constructed by PCR amplifying the *aacC1* gene (Gm^R^), from pSEVA611 plasmid^28^, to which 33-bp, encoding the AANDENYALAA protease degradation tag was added to the 3-end using primers Gm-kpnI-F and GmLAA-hindIII-R (Table S1). The ~0.5-kb PCR DNA was digested with KpnI and HindIII and used to replace the ~0.7-kb fragment containing the *cat*LVA resistance marker from pZS4int-rhl-catLVA^22^ generating the pZS4int-rhIL-GmLAA plasmid. Thus, the “Genomic insert” parental strain was constructed by integrating the cassette T0-Sp^R^-*rhlR←P*_*lacIq*_-*P*_*rhl*_→*Gm*^R^_LAA_-T_1_ into the λ attachment site (*attB*) of *E. coli* JC1080 with the helper plasmid pLDR8, that bears the lambda integrase, as described^29^. The pZS4int-rhIL-GFP plasmid was assembled by extracting the *gfp* gene from pSEVA241-*P*_*rhl*_→*gfp* vector (lab collection) upon digestion with KpnI and HindIII. Then, the 0.7-kb DNA fragment was cloned into the KpnI and HindIII sites of pZS4int-rhIL-GmLAA replacing the Gm^R^_LAA_ cassette for a GFP fluorescent reporter.

Then, the quorum sensing reporter “biosensor” strain was constructed by introducing into the λ attachment site (*attB*) of *E. coli* JC1080 the cassette T_0_-Sp^R^-*rhlR←P*_*lacIq*_-*P*_*rhl*_→,*gfp*-T_1_ as described above. The plasmid pZS*2R-GFP-rhlI (ori-SC101*, *P*_*R*_→*gfp-rhlI*, Km^R^) was obtained from^22^. The pZS*2R-mCherry (ori-SC101*, *P*_*R*_→*;gfp-rhlI*, Km^R^) plasmid was constructed by amplifying *mcherry* from pSEVA237R (4)(9) using primers mCherry-kpnI-F and mCherry-xbaI-R (Table S1). The PCR-amplified fragment was flanked with KpnI and XbaI restriction sites and cloned into the corresponding sites of the ~3.5-kb pZS*2R-GFP-rhlI plasmid backbone, thereby swapping the *gfp-rhlI* cassette for the *mcherry* gene.

### GFP expression assay to estimate QS signal concentration

To estimate the concentration of the quorum sensing signal produced (C4-HSL), in different experimental conditions we used the reporter “biosensor” bacteria. We grew the “biosensor” strain overnight, adjusted the OD_600_ to 0.1 and grew aerobically for 2.5h at 30°C. Then, the culture was aliquoted and purified supernatant from the experimental sample was added. We used known concentrations (0, 0.1μM, 1μM, 10μM, and 100μM) of the commercial N-butyryl-L-homoserine lactone (C4-HSL) to obtain a calibration curve. We grew these preparations aerobically for 3h, laid out 200 μl of cultures in quadruplicates in a 96-well black/clear bottom microtiter plate (Sigma-Aldrich, MO, USA) and measured OD_600_ and GFP fluorescence (ex: 488nm; em: 520nm; cutoff: 495nm) in a SpectraMax M2e microplate reader (Molecular Devices, CA, USA). Supernatants were obtained from 200μl of overnight liquid cultures prepared following the “Accumulation of PG and stress” protocol, centrifuged twice at room temperature to remove cells and used directly to induce growing cells of the reporter strain. Comparing observed supernatant fluorescence to the calibration curve approximated QS molecule quantities.

### Engineering of initial conditions

Initial populations for the “accumulation of PG and stress” protocol and others were prepared by mixing cooperators and cheaters at a defined population density (10^4^ cells/well for high initial density experiments and 1-10 cell/well for low initial density experiments) and cooperator frequency. Overnight cultures of producers and nonproducers were washed twice with PBS by centrifugation for 15min at 3800rpm and room temperature. Then, OD_600_ was adjusted to 0.15. We assembled populations at the desired P frequency in a fixed final volume (2.5ml), which was then serially diluted to the required cell density. This dilution was done in large volumes of medium (20ml) and applying low dilution factor (¼) each step to minimize the introduction of error in strain frequencies. Initial dilution steps were performed in PBS and the final 3 steps were performed in LB with Km, Sp. The robustness of this procedure is shown in Fig. S5.

### Invasion of nonproducers

We prepared washed cultures of producers and nonproducers as described above. After adjusting OD_600_, to 0.15 we mixed both strains at the indicated frequency. Then, we inoculated three replica 50ml Erlenmeyer flasks with 5ml of LB, Km, Sp. After 24h, we reseeded a new flask with a 1/100 dilution and fresh medium. In order to estimate producer frequency in grown cultures, cells were 1/10 serially diluted in PBS using a total of 10ml of medium, plated onto LB agar plates, and colonies counted after 24h at 30°C. We followed this process for 4 consecutive days.

### “Accumulation ofPG and stress”

Populations with a given initial cell density and cooperator frequency were prepared, distributed into a 96-multiwell plate and incubated for 15.5h (T_1_). Then, 1/10 of each well was transferred into a new 96-multiwell plate with LB, Km, Sp and the specified gm concentration. This plate is again incubated for 8.5h (T_2_). At the end of T_2_ we plated whole well content on LB agar plates and counted CFU after 48hrs at 30°C.

### Antibiotic sensitivity

Overnight cultures were reseeded and grown for 4h at 30°Cto reach exponential phase. Aliquots of this culture containing ~10^6^ cells were resuspended into a 96-multiwell plate with LB and a given dose of gm, incubated for 2h or 4h, plated and counted. For experiments with the nP strain, Km and Sp were added to the LB medium. For experiments with cells in stationary phase, overnight cultures were used directly in the initial inoculation. We used additional wells without antibiotic in parallel to obtain a reference population size. We express the sensitivity to the antibiotic as “fraction surviving” (population size after exposure to antibiotic / reference population size without antibiotic).

### Effect of synthetic quorum-sensing on gentamicin tolerance

Overnight cultures of nonproducers were resuspended into LB with Km, Sp and the indicated concentrations of synthetic *quorum-sensing* molecule (C4-HSL). Then, we proceeded with the antibiotic sensitivity assay as described above.

### Mutation rate

We generated replica populations of the nP strain and initial density of ~1 cell/well. We distributed these cultures in 96-multiwell plates and allowed them to grow for 15.5h (T_1_). We plated the whole well content on LB agar plates with the specified gm dosage and counted viable cells. From the distribution of gmR CFU observed in a set of replica populations the mutation rate was estimated with a maximum likelihood method as described in30, using the online application “FALCOR: Fluctuation Analysis Calculator” (http://www.keshavsingh.org/protocols/FALCOR.html)

